# Molecular basis of SARS-CoV-2 Omicron variant evasion from shared neutralizing antibody response

**DOI:** 10.1101/2022.10.24.513517

**Authors:** Anamika Patel, Sanjeev Kumar, Lilin Lai, Chennareddy Chakravarthy, Rajesh Valanparambil, Elluri Seetharami Reddy, Kamalvishnu Gottimukkala, Prashant Bajpai, Dinesh Ravindra Raju, Venkata Viswanadh Edara, Meredith E. Davis-Gardner, Susanne Linderman, Kritika Dixit, Pragati Sharma, Grace Mantus, Narayanaiah Cheedarla, Hans P. Verkerke, Filipp Frank, Andrew S. Neish, John D. Roback, Carl W. Davis, Jens Wrammert, Rafi Ahmed, Mehul S. Suthar, Amit Sharma, Kaja Murali-Krishna, Anmol Chandele, Eric A. Ortlund

## Abstract

A detailed understanding of the molecular features of the neutralizing epitopes developed by viral escape mutants is important for predicting and developing vaccines or therapeutic antibodies against continuously emerging SARS-CoV-2 variants. Here, we report three human monoclonal antibodies (mAbs) generated from COVID-19 recovered individuals during first wave of pandemic in India. These mAbs had publicly shared near germline gene usage and potently neutralized Alpha and Delta, but poorly neutralized Beta and completely failed to neutralize Omicron BA.1 SARS-CoV-2 variants. Structural analysis of these three mAbs in complex with trimeric spike protein showed that all three mAbs are involved in bivalent spike binding with two mAbs targeting class-1 and one targeting class-4 Receptor Binding Domain (RBD) epitope. Comparison of immunogenetic makeup, structure, and function of these three mAbs with our recently reported class-3 RBD binding mAb that potently neutralized all SARS-CoV-2 variants revealed precise antibody footprint, specific molecular interactions associated with the most potent multi-variant binding / neutralization efficacy. This knowledge has timely significance for understanding how a combination of certain mutations affect the binding or neutralization of an antibody and thus have implications for predicting structural features of emerging SARS-CoV-2 escape variants and to develop vaccines or therapeutic antibodies against these.

## Introduction

SARS-CoV-2 Omicron subvariants are continuously emerging and escaping therapeutic monoclonal antibodies (mAbs) and vaccines (1–3). Mutations acquired in the spike protein of SARS-CoV-2 variants, a target for neutralizing antibodies (nAbs), are primarily responsible for this immune escape (1, 4). Identifying nAbs / non-nAbs to these variants and determining their prevalence in human population allows us to understand the shared mechanisms of immune protection among diverse populations (5, 6). Since the emergence of COVID-19, >11,000 SARS-CoV-2 mAbs have been identified (7). Among these, nAbs encoded by human antibody heavy chain variable germline genes such as IGHV3-53/3-66, IGHV1-58, IGHV3-30 and IGHV1-69 are commonly observed in many individuals across the globe (7). These related rearrangements, known as a public antibody response, suggest a shared immune response with a similar genetic makeup and modes of antigen recognition that has been found in large number of individuals infected with influenza, dengue, malaria, HIV and SARS-CoV-2 (5, 6, 8–13). Mapping the immunogenetic makeup, structure, and function of these public clonotypes allows us to better understand how certain mutations affect the binding of an antibody and thus potentially expedite antibody re-purposing for emerging variants. It is established that SARS-CoV-2 variants bearing K417N/N501Y mutations evade IGHV3-53/3-66 RBD mAbs (5, 13). These antibodies are primarily encoded by near germline sequences and are commonly found in populations residing in distinct geographical regions (5, 12, 13). However, SARS-CoV-2 variant evasion from the IGHV3-30 shared antibody response is unclear.

We recently published a panel of 92 RBD-binding monoclonal antibodies (mAbs) isolated from five individuals infected with the ancestral SARS-CoV2 strain in India and identified a potent class-3 broad-spectrum antibody capable of neutralizing all highly evasive Omicron variants (14, 15). Here, we focused on three mAbs that potently neutralize the ancestral WA.1 strain, but differentially neutralize SARS-CoV-2 variants for further characterization. The immunogenetic analysis confirms that all three mAbs were encoded by IGHV3-53/66 and IGHV3-30 genes and were publicly shared (14). While the Cryo-EM structure of all three mAbs showed bivalent spike binding, two mAbs (002-02 and 034-32) targeted the class-1 RBD epitope whereas mAb 002-13 targeted a relatively conserved class-4 epitope. Detailed look of molecular interactions at each mAb’s epitope-paratope surface allowed us to predict how mutations of certain residues in key variants of concern (VOCs) might impact antibody functionality and their role in immune evasion.

## Results

### Identification and characterization of shared human mAbs to SARS-CoV-2

In this study, we have selected three out of 92 previously identified RBD-specific mAbs for further characterization (14). These three mAbs, referred to as 002-13, 002-02, and 034-32 have heavy chain VJ pairings encoded by IGHV3-30, IGHD2-8, IGHJ4; IGHV3-66, IGHD4-17, IGHJ4 and IGHV3-53, IGHD1-1, IGHJ6 immunoglobulin genes, respectively, whereas their light chain VJ pairings were encoded by IGLV6-57, IGLJ2; IGK3-20, IGKJ4 and IGK1-9, IGKJ3 genes, respectively (**Figure 1A**). Genetic analysis of these three mAbs showed that heavy chain variable (V)-genes of all three mAbs were encoded by a shared public antibody response (**Figure 1B, 1C, 1D, 1E and S1**) as documented in the CoV-AbDab database of all RBD-specific mAbs (n=6520) isolated from SARS-CoV-2 infected/vaccinated individuals (7). Interestingly, the antibody gene IGHV3-30, IGHJ4 of 002-13 mAb is the most frequent VJ pairing used by SARS-CoV-2 RBD mAbs (**Figure 1D**). Heavy chain V-gene IGHV3-30 of mAb 002-13 is the second most frequently used IGHV gene among all RBD mAbs (**Figure 1B**). Interestingly, 002-13 like shared mAbs exhibit the presence of a conserved CxGGxC motif in their 22-residue long complementarity determining region (CDR) H3 (CDRH3) (**Figure S1A**) encoded by a IGHD2-8 gene (7). The IGHV genes of 002-02 (IGHV3-66) and 034-32 (IGHV3-53) have already been described earlier in detail as a shared clonotype antibody response that shows the characteristic motifs of NY and SGGS in their CDRH1 and CDRH2 regions, respectively, preferred IGHD4- 17 gene and a short CDRH3 length of 9 – 12 amino acids with high sequence diversity (5, 12, 13) (**Figure S1B and S1C**) (7).

**Figure 1:**
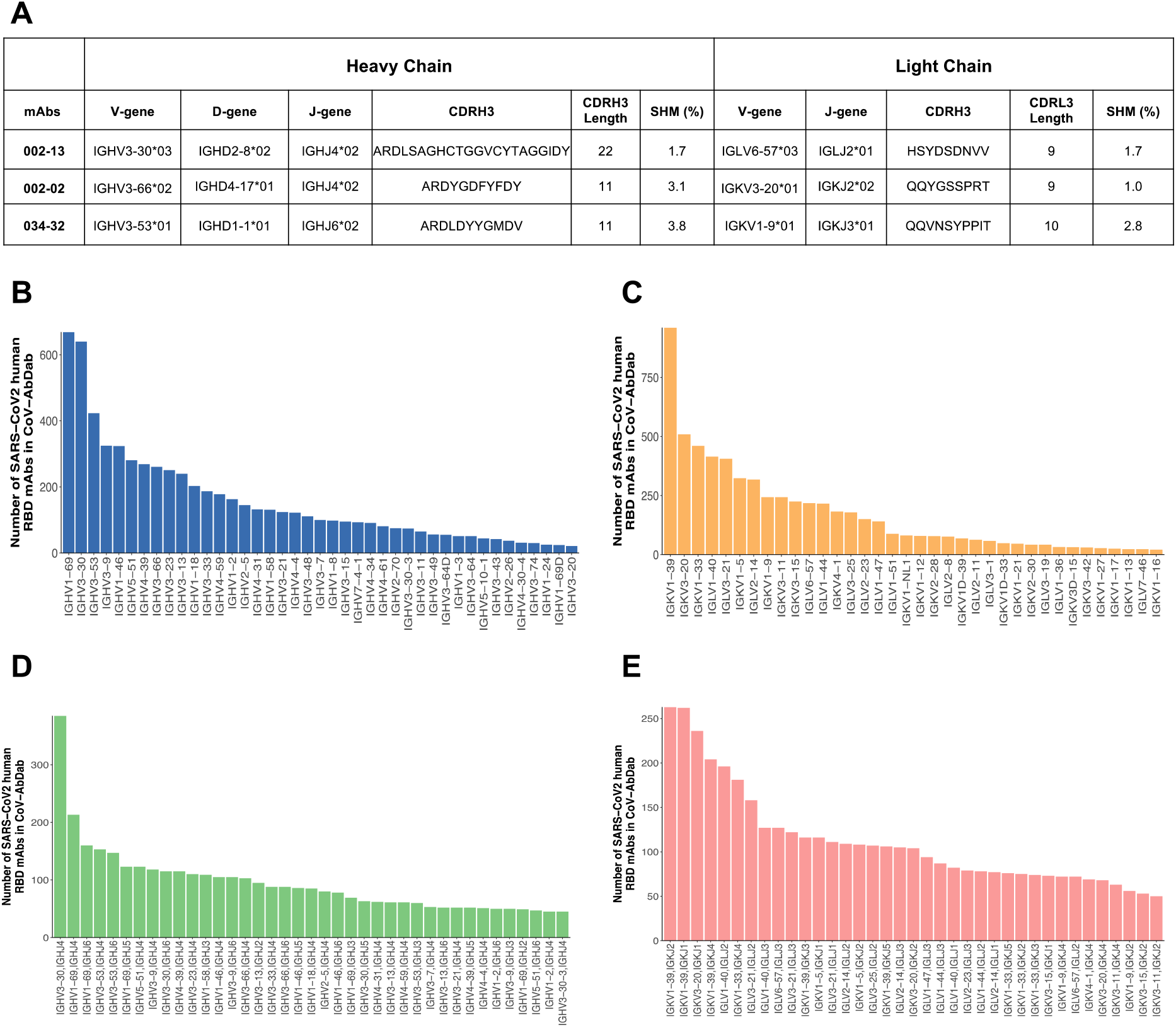
Genetic information of SARS-CoV-2 RBD specific shared mAbs. (**A**) Immunogenetic information of the three SARS-CoV-2 mAbs. (**B**) Heavy chain variable gene distribution of SARS-CoV-2 RBD-specific human mAbs (N=6520) documented in CoV-AbDab dataset. (**C**) Light chain variable gene distribution of SARS-CoV-2 RBD-specific human mAbs documented in CoV-AbDab dataset. (**D**) Heavy chain VJ-gene bar plot of SARS-CoV-2 RBD-specific human mAbs documented in CoV-AbDab dataset. (**E**) Light chain VJ-gene bar plot of SARS-CoV-2 RBD-specific human mAbs documented in CoV-AbDab dataset.

Next, we revealed that all three mAbs strongly bind spike protein with Kd values in low nM to pM range, by both BLI (**Figure S2**) and Mesoscale binding assay (Mesoscale Discovery) (**Figure 2A**). Additionally, in agreement with binding data they all potently neutralize the ancestral WA.1 live virus in a focus-reduction neutralization mNeonGreen (FRNT-mNG) assay (**Figure 2B and 2C**) (14, 15). Taken together, these results confirm high binding affinity and potent neutralizing capacity of all three shared mAbs against the SARS-CoV-2 WA.1 strain.

**Figure 2:**
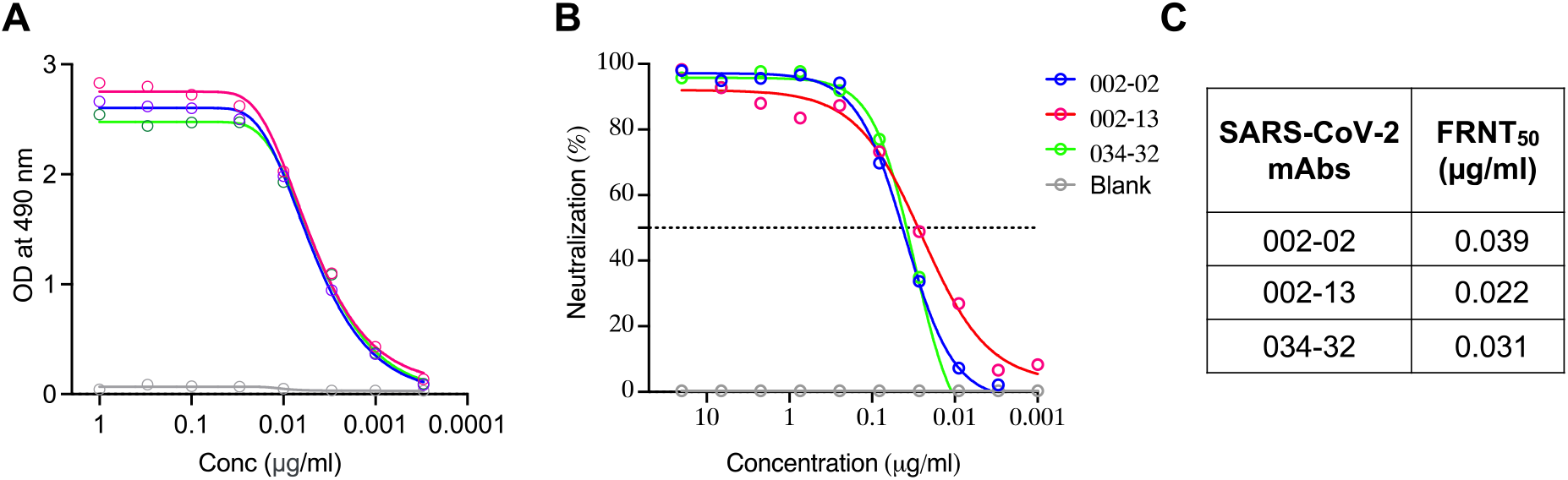
Binding, neutralization and affinity analysis of selected mAbs towards the WA.1 strain. **(A)** Three SARS-CoV-2 mAbs were tested for binding to the WA.1 RBD protein. **(B)** Live virus neutralization curves of the three mAbs against live WA.1 SARS-CoV-2. Neutralization was determined on using a focus reduction neutralization mNeonGreen (FRNT-mNG) assay on Vero cells. **(C)** 50% focus reduction neutralization titres (FRNT-mNG50) for the three SARS-CoV-2 mAbs against WA.1 are shown.

### Epitope mapping of mAbs 002-02, 002-13 and 034-32

To delineate the molecular determinants conferring epitope recognition and to understand the mechanism of their potent neutralization against WA.1 strain, we solved the Cryo-EM structures of WA.1 spike-6P (Spike-hexapro) in complex with each of the three mAbs (002-13, 002-02 and 034-32) in their native full-length IgG form (**Figure 3 and 4**). The structures show bivalent binding modes for all three mAbs, revealing two distinct neutralization mechanisms (**Figure S4**). Below we summarize our observations.

**Figure 3.**
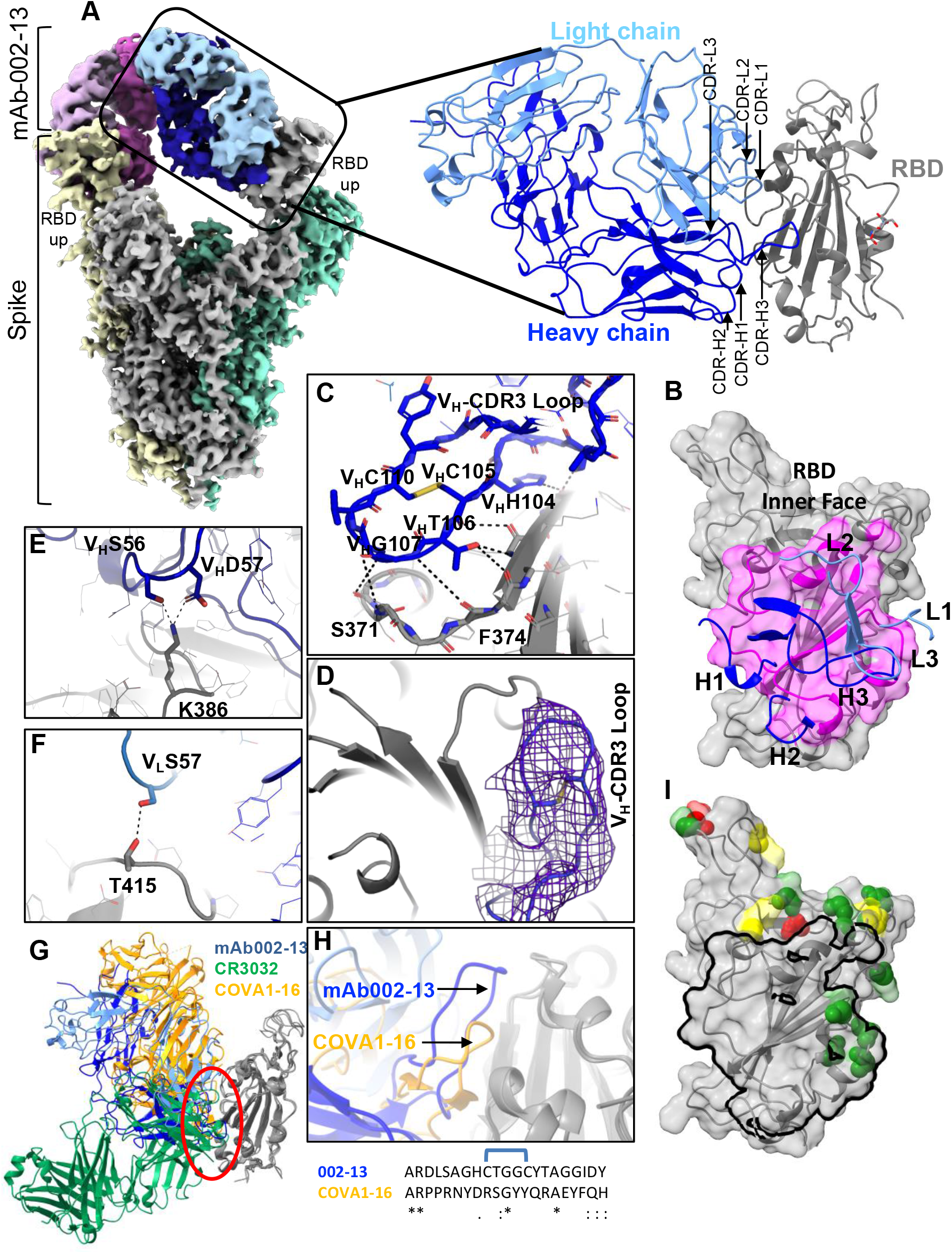
Cryo-EM structure of 002-13 in complex with WA.1 Spike trimer explains its broad neutralization activity. **(A)** Cryo-EM structure of WA.1 Spike-6P trimer in complex with mAb 002-13. Overall density map at contour level of 5.4 *σ* showing the antibody binding in the RBD “up” conformation. Each Spike protomer is shown in gray, yellow or green; light and heavy chains of each Fab region are shown in blue/ magenta and light blue/ pink, respectively. A model for one complex between Fab and RBD is shown to the right. The positions of all Fab complementarity-determining region (CDR) regions are labelled. **(B)** Surface representation of RBD with relative positions of all CDR loops. The mapped epitope surface in the RBD is highlighted in magenta. **(C, E, F)** Interaction details at the 002-13-RBD interface. **(D)** Heavy chain CDR3 loop in density map. **(G)** Comparison of 002-13 binding mode with other Class-4 mAbs. **(H)** Zoom in view comparing the heavy chain CDR3 loop positions of 002-13 vs COVA1-16. CDR3 amino acid sequence of 002-13 and COVA1-16 is shown below. **(I)** Locations of beta (yellow), delta (red) and omicron (green) mutations on the RBD relative to the 002-13 epitope site (black outline).

**Figure 4:**
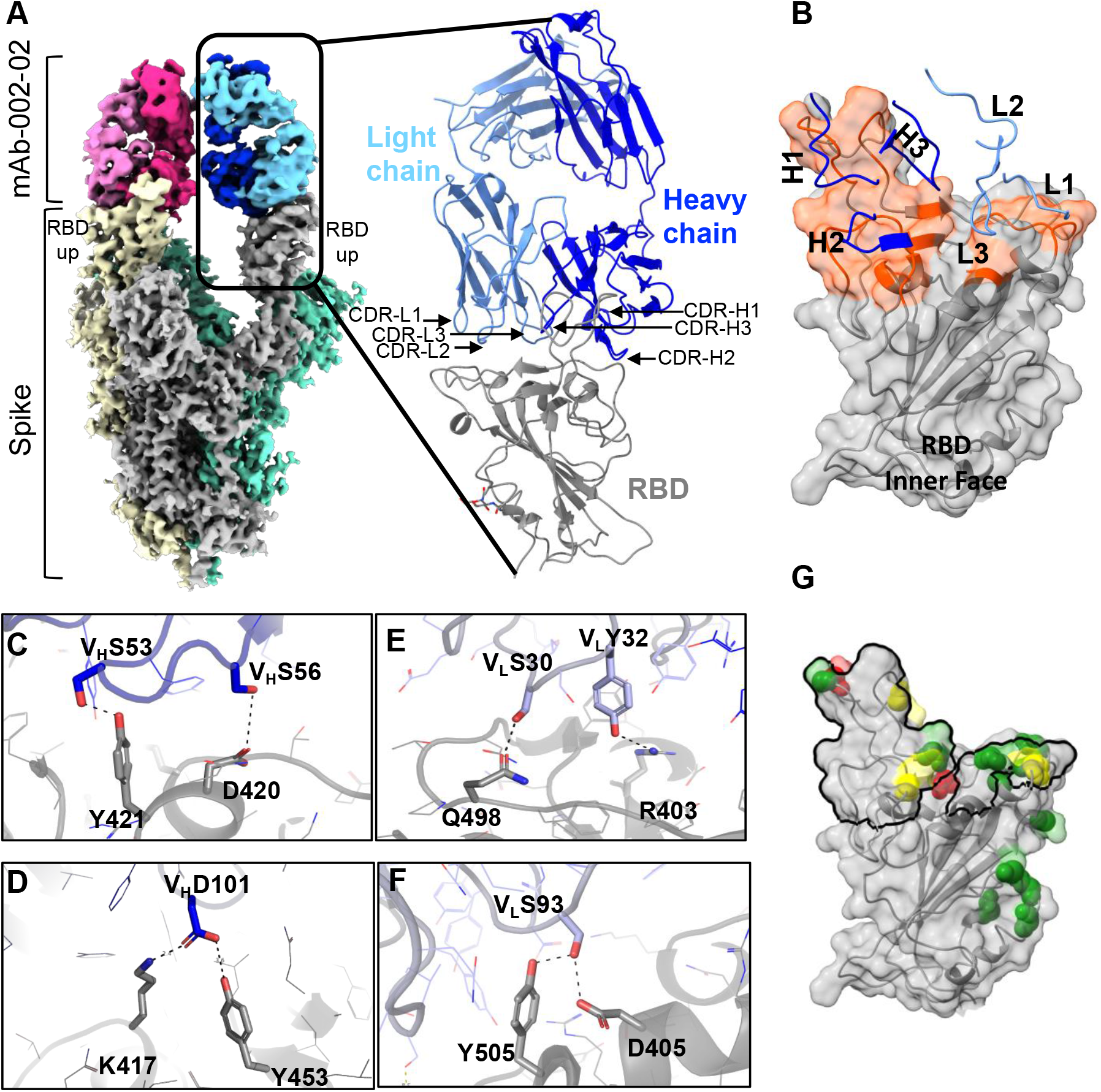
Cryo-EM structure of 002-02 in complex with WA.1 spike trimer. **(A)** Cryo-EM structure of WA.1 spike-6P trimer in complex with mAb 002-02. Overall density map at contour level of 3.6 *σ* showing the antibody binding two RBDs in the “up” conformation. Each protomer of Spike is shown in gray, yellow or green; the light and heavy chains of each Fab region are shown in blue/ magenta and light blue/ pink, respectively. A model one Fab-RBD complex is shown to the right and the positions of all Fab CDR regions are labelled. **(B)** Surface representation of the RBD showing the relative positions of all CDR loops. The mapped epitope surface in the RBD is highlighted in orange. **(C-F)** Interaction details of the 002-02-RBD interface. **(G)** locations of Beta (yellow), Delta (red) and Omicron (green) mutations on RBD relative to the 002-02 epitope site (black outline).

**mAb 002-13:** The Cryo-EM structure of 002-13 in complex with WA.1 spike-6P (**Figure 3 and S3**) resolved at 3.8 Å global resolution revealed a conserved epitope on the inner face of the RBD, aligning with RBD-7/class-4 epitopes only accessible in up conformation. There is clear intra-spike bivalent binding, where each Fab region of the full-length IgG recognizes adjacent RBDs in a spike trimer (**Figure 3A and S4**). 002-13 mAb, belongs to the public clonotype encoded by IGHV3-30 and IGLV6-57 that have not been structurally characterized before. Notably, the 22-residue long CDRH3 region encoded by IGHD2-8 gene of 002-13 mAb contains a CxGGxC motif which is shared by other 81 RBD-specific mAbs (**Figure S1A**) documented in CoV-AbDab database (7). Like other class-4 antibodies, 002-13 RBD binding is dominated by the heavy chain contributing ~76% of total interaction with a total buried surface area of ~887 Å^2^ (**Figure 3B**) (16, 17). Most of the heavy chain interactions are mediated through the CDR3 region that forms a foot-like loop, stabilized by an intra-loop disulfide bond between residues C105 and C110 of CxGGxC motif (**Figure 3C and 3D**). We observe that multiple interactions involving the residues in RBD region S371- C379 and the heavy chain CDR3 loop are responsible for epitope recognition (**Figure 3C**). Heavy chain CDR2 residues D57 and S56 engage RBD residue K386 through a salt bridge and hydrogen bond (**Figure 3E**). The light chain of 002-13 contributes minimally to RBD binding, only the CDR2 loop of light chain comes into RBD proximity to make a hydrogen bond with the side chain of RBD residue T415 (**Figure 3F**). Although 002-13 binds outside the Receptor Binding Motif (RBM) surface, it can sterically block ACE2 binding through its light chain orientation as previously observed in ACE2 competition profiling (14).

Structural comparison of 002-13 with another class-4 monoclonal antibody (COVA1-16 and CR3032) shows a distinct binding pose for 002-13 (**Figure 3G**), additionally, a unique small side chain-containing sequence in CxGGxC motif of the heavy chain CRD3 loop allows it to go much deeper into the RBD pocket facilitating extensive interactions in this region compared to other class-4 mAbs (**Figure 3H**).

We then marked key mutations present in Beta (yellow), Delta (red) and Omicron (green) variants within the 002-13 epitope surface and observed that while all VOC except for Omicron carry no mutations, Omicron carries three mutations (S371L, S373P and S375F) within the 002-13 epitope (**Figure 3I**). This suggests that while the binding and neutralization of 002-13 mAb will be preserved for most SARS-CoV-2 variants, it might be impacted towards Omicron as these mutations are known to induce a local conformational change in the Omicron RBD structure and thus, could exclusively evade Omicron (18, 19).

**mAb 002-02 and 034-32:** Antibodies 002-02 and 034-32 were isolated from two different individuals and are encoded by public clonotype genes IGHV3-53/3-66 (**Figure 1A**). They both show very similar properties with high binding specificity towards SARS-CoV-2 RBD, effectively compete ACE2 and potently neutralize WA.1 (14) (**Figure 2**). To define the details of epitope recognition, we determined the Cryo-EM structure of each 002-02 (**Figure 4 and S5**) and 034-32 (**Figure S6 and S7**) in complex with WA.1 Spike- 6P at a resolution of 3.8 and 4.3 Å, respectively. For both complexes, we observe intra-spike bivalent binding, where each Fab region of IgG binds two neighboring RBDs in the spike trimer in the “up” conformation. The RBD that does not engage in binding Fab remains in the “down” conformation (**Figure 4A)**. Both mAb structures recognize epitopes in the top RBD pocket that aligns with the RBM surface suggesting direct ACE2 competition and based on this they are classified as RBD-2/class-1 antibodies. Since 002-02 and 034-32 recognize the RBD in a very similar manner, we focus our structural analysis on mAb-RBD recognition in the locally refined map for 002-02.

While all CDR loops are involved in epitope recognition (**Figure 4A and 4B**), most RBD contacts are dominated by the heavy chain, contributing ~70% of the total of 1058 Å^2^ of buried surface between mAb and RBD (**Figure 4B**). Primary interactions in the heavy chain are mediated by CDR1 and CDR2 regions. Most mAbs that belong to class-1 antibodies are encoded by public clonotype genes IGHV3-53/IGHV3-66 (5, 13). The common features among these mAbs include a conserved NY and SGGS motif in CDR1 and CDR2 regions, respectively, that contribute significantly toward RBD binding (12). We also observed a network of hydrogen bonds with the RBD through the CDR2 SGGS motif. The side chain of S53 and S56 in the CDR2 heavy chain engages in a hydrogen bond with side chains of Y421 and D420 in RBD, respectively (**Figure 4C**). However, the CDR3 loop heavy chain residues in this mAb class varies more. In 002-02, the heavy chain CDR3 residue D101 forms a hydrogen bond with K417 and Y453 in the RBD (**Figure 4D**). In the light chain, CDR1 and CDR3 make some contact with the inner left side of the RBD. The S30 and Y32 residue in the CDR1 region of the light chain makes a hydrogen bond with Q498 and R403 in RBD, respectively (**Figure 4E**). Also, the S93 in the CDR3 region of the light chain interacts with Y505 and D405 (**Figure 4F**).

Like 002-13, we also mapped mutations found in Beta (yellow), Delta (red) and Omicron (green) variants onto the 002-02 / 034-32 epitope (**Figure 4G**). While Delta carries no mutation within 002-02 / 034-32 epitope surface, three of the Beta mutations (K417N, E484K, N501Y) fell within its epitope, suggesting no variation in binding and neutralization for Delta but weakened binding and neutralization for Beta. However, six Omicron mutations (K417N, S477N, Q493R, G496S, Q498R and N501Y) lied within the 002-02/ 034-32 epitope surface and predicted to evade Omicron binding and neutralization. Collectively, based on these observations both 002-02 and 034-32 mAbs will be less or ineffective towards both Beta and Omicron variants.

### Assessing binding and neutralization breadth towards SARS-CoV-2 variants

To link the paratope mutation landscape in VOC to the antibody function, we tested binding and neutralization of these three mAbs against SARS-CoV-2 variants. In agreement with the structure-based prediction, the binding of 002-13 (class-4 antibody) remained unaffected towards Alpha, Beta and Delta variants as these variants contain no mutations within the 002-13 epitope and showed moderately reduced (~2.7-fold) binding to Omicron (**Figure 3I, Figure 5A and 5G**). In agreement with binding data, the neutralization potency of 002-13 remained unperturbed in Alpha, Beta, Delta and showed no observable neutralization of the Omicron virus (**Figure 5E and 5G**). Along that line, binding of 002-02 and 034-32 (class-1 antibodies) retained for Alpha and Delta variants to the same affinity as of WA.1, showed 3-fold and 150-fold reduced affinity to Beta, respectively, and no observable binding to Omicron (**Figure 5B, 5C and 5G**). Following this tread both 002-02 and 034-32 neutralize Alpha and Delta variants with the same potency as WA.1, showed 4-fold and 17-fold reduced potency to Beta, respectively and complete loss of neutralization to Omicron (**Figure 5E, 5F and 5G**). This is further supported by the fact that unique K417N mutation (present in Beta but not in Delta) would result in a loss of a hydrogen bond with D101 in heavy chain CDR3 (**Figure 4D**) and subsequent Beta-variant specific loss of binding and neutralization for 002-02 and 34-32. This was also confirmed by a 2-fold decrease in the calculated ddG value of - 46.23 +/- 10.5 kcal/mol based on molecular mechanics/ Poisson-Boltzmann surface area (MM/PBSA) free energy for the single K417N mutant in 002-02-spike structure compared to the WA.1 ddG value of -82.62 +/- 9.57 (**Figure S8**).

**Figure 5:**
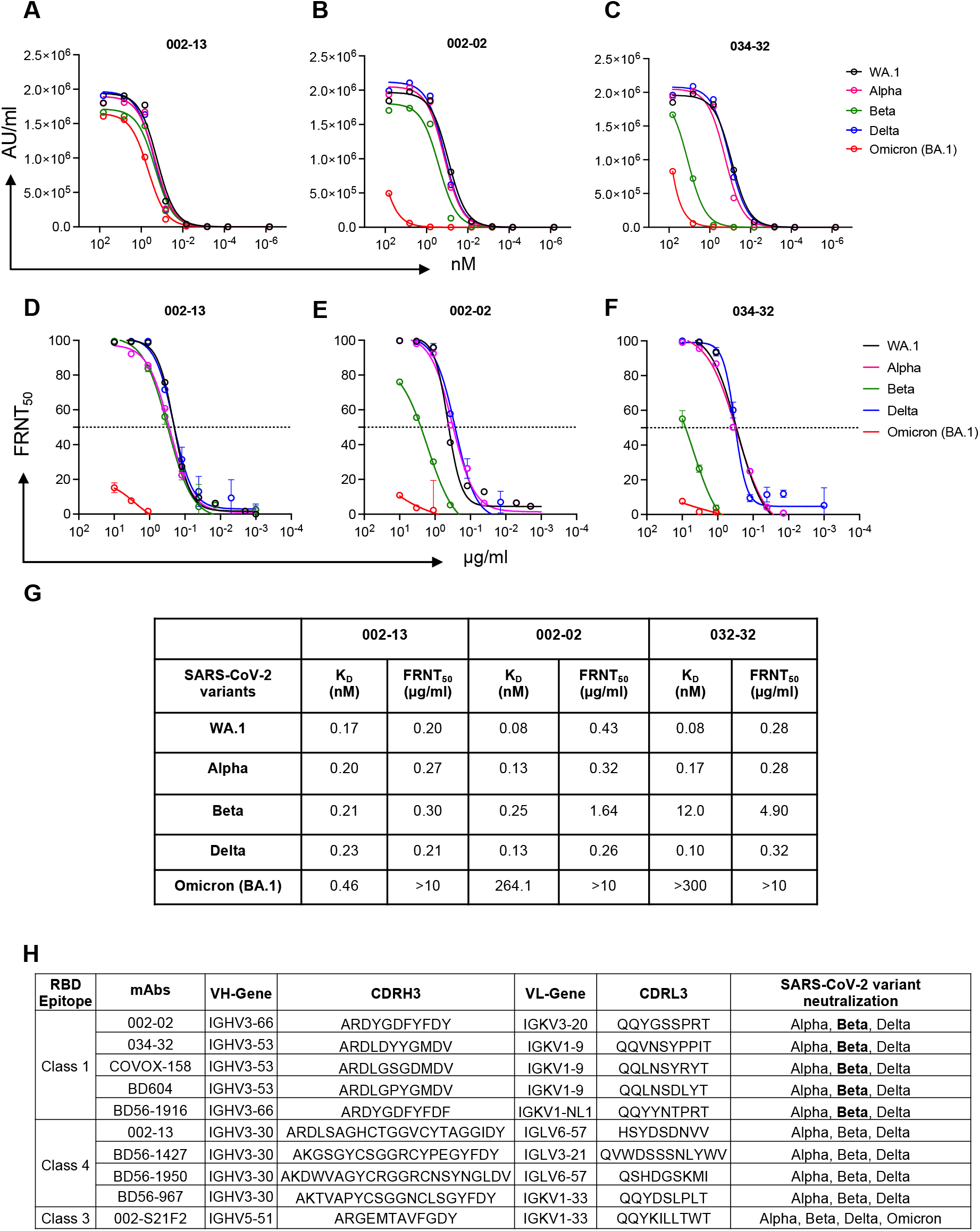
Binding affinity and neutralization analysis of selected mAbs against SARS-CoV-2 variants. **(A-C)** Three potent neutralizing mAbs were tested for binding to spike proteins of SARS-CoV-2 WA.1, Alpha, Beta, Delta and Omicron (BA.1) variants of concern (VOCs). Curves shown are the best fit one-site binding curves calculated by Prism 9.0 **(D-F)** Live virus neutralization curves and FRNT50 values of three potent mAbs for WA.1, Alpha, Beta, Delta and Omicron (BA.1) SARS-CoV-2 VOCs are shown. Neutralization was determined on Vero-TMPRSS2 cells using a focus reduction neutralization assay. **(G)** Table summarizing the dissociation constant (K_D_) and neutralization potency of mAbs against SARS-CoV-2 variants. **(H)** Comparison of three mAbs with other similar RBD epitope class recognizing mAbs and their reported neutralization to SARS-CoV-2 variants (variants in Bold show reduced potency) (7).

Altogether, this data catalogues the epitope class-specific antibody susceptibility towards existing SARS-CoV-2 variant and can inform their action on a newly emerging variant.

## Discussion

Understanding how SARS-CoV-2 mAbs achieve broad neutralization or are rendered ineffective by viral mutations provides insight not only about natural immunity, but is critical to develop broadly effective therapeutic mAbs and guide vaccine design (5, 12, 20–22). Moreover, defining antibody-antigen interactions is critical for the rapid re-evaluation of existing antibody-based therapeutics towards continuously emerging SARS-CoV-2 variants. This, overlaid with the immuno-genetic makeup of the antibodies shared by large population further informs our understanding of the public immune response and their antigenic drift from variants. For example, certain antibody responses are repeatedly shared among large number of individuals regardless of their genetic origins, as has been observed previously during different pathogen infections including influenza, dengue, HIV and Malaria (8–11). With SARS-CoV-2, these are encoded by IGHV3-53/66, IGHV1-58, IGHV3-30 and IGHV1-69 which are found both following natural infection and post-vaccination (5, 6, 12). Such information can be collectively used to fine tune the immune response focused on broad and potent neutralizing epitopes through antigen design for a universal vaccine (20, 22). Recently, based on the information from shared public clonotypes of HIV-1 bnAbs, a V2-apex region specific immunogen has been successfully designed (23).

We recently reported the isolation of 92 SARS-CoV-2 RBD-specific mAbs from COVID-19 recovered individuals from India during the first wave of the pandemic and identified a broadly neutralizing class-3 antibody (002-S21F2), capable of neutralizing all omicron subvariants (14). Out of 92, three SARS-CoV-2 nAbs (002-13, 002-02 and 034-32) characterized in this study, belong to shared public antibody responses. Sequence analysis of 6520 published SARS CoV-2 RBD specific mAbs define 002-13 as a public clonotype encoded by IGHV3-30, IGHJ4 genes with >80% of these exhibiting IGHD2-8 gene usage and presence of CxGGxC motif in their CDRH3 region that have not been structurally characterized (7). While the other two mAbs, 002-02 and 034-32 mAbs are encoded by shared IGHV3-53/3-66 antibody genes as previously shown by others (5, 12, 13). The Cryo-EM structures for these three mAbs in complex with trimeric spike protein show class-4 epitope recognition by 002-13 and class-1 epitope recognition by 002-02 and 034-32. The structures further allowed us to define their epitope-paratope interfaces in detail in relation to the locations of SARS-CoV-2 variants mutations to predict viral immune escape. While there was no observable difference in the antibody functionality for variants containing mutations that lie outside the mapped epitope surface of a particular antibody, there was a remarkable drop in binding affinity, and neutralization of the antibody when the mutations mapped to the antibody footprint. Most broad neutralizing antibodies recognize all variants antigen that either carry no mutations within their epitopes or the mutations in epitope region are favored by mAb specific molecular interactions as we observed for class-3 mAb 002-S21F2 (14). Here, we show all three mAbs potently neutralized the ancestral WA.1 strain, but differentially neutralize other variants, primarily due to the presence of evading mutations present in their epitope antigenic sites, similar to the other well characterized mAbs recognizing the same epitope classes (**Figure 5H**) (7). Major mutations responsible for Beta evasion are K417N, E484A for 002-02 and 034-32 mAbs, also observed previously for IGHV3-53/3- 66 shared antibody responses (5, 12). Omicron, which contains six epitope mutations (K417N, S477N, Q493R, G496S, Q498R and N501Y) within 002-02/034-32 and three mutations (S371L, S373P and S375F) within 002-13 binding site, would collectively lead to major immuno-escape, especially as some mutation residues participate in direct interaction with mAb. Although 002-13 showed only moderate reduction in binding affinity, it showed no neutralization towards Omicron suggesting additional factors might play a role in 002-13 specific Omicron escape. One explanation could be that Omicron mutations that favor Spike “up” conformation would likely promote ACE2 interaction and reduce 002-13 mAb competition (24). Our findings suggest that immune pressures exerted by the shared antibody response to SARS-CoV-2 are likely to cause evolution variants with mutations in the class-4 antibody epitope residues S371, S373 and S375. These mutations must be tracked to find effective solutions to combat emerging variants. Further, the structure guided prediction made for three SARS-CoV-2 shared nAbs that potently neutralized the WA.1 strain holds true towards the functional efficacy of these mAbs against SARS-CoV-2 variants, including Omicron.

In summary, this study vastly improves our understanding of how Omicron escaped from shared antibody responses to SARS-CoV-2 elicited during the natural infection and has implications towards concepts for fast-tracking effective broad range therapeutics against continuously emerging SARS-CoV-2 variants.

## Materials and Methods

### SARS-CoV-2 RBD-specific ELISA binding assays

The recombinant SARS-CoV-2 RBD gene was cloned, expressed, purified and ELISAs were performed as previously described (14, 15, 25). Briefly, purified RBD was coated on 96- well MaxiSorp plates (Thermo Fisher, #439454) at a concentration of 1 μg/mL in phosphate-buffered saline (PBS) at 4°C overnight. The plates were washed with PBS containing 0.05% Tween-20. Three-fold serially diluted purified mAb was added and incubated at room temperature for 1 hr. Plates were washed and the SARS-CoV-2 RBD specific IgG signal was detected by incubating with horseradish peroxidase (HRP) conjugated - anti-human IgG (Jackson ImmunoResearch Labs, #109-036-098). Plates were then washed thoroughly and developed with o-phenylenediamine (OPD) substrate (Sigma, #P8787) in 0.05M phosphate-citrate buffer (Sigma, #P4809) pH 5.0, containing 0.012% hydrogen peroxide (Fisher Scientific, #18755). Absorbance was measured at 490 nm.

### Live SARS-CoV-2 neutralization assay

Neutralization titers to SARS-CoV-2 were determined based on either a focus-reduction neutralization mNeonGreen (FRNT-mNG) assay on Vero cells or FRNT assays based on Vero TMPRSS2 cells as previously described (14, 15). Briefly, 100 pfu of SARS-CoV-2 (2019-nCoV/USA_WA1/2020), Alpha, Beta, Gamma, Delta and Omicron variants were used on Vero TMPRSS2 cells. Purified monoclonal was serially diluted three-fold in duplicate starting at 10 μg/ml in a 96-well round-bottom plate and incubated for 1 h at 37°C. This antibody-virus mixture was transferred into the wells seeded with Vero- TMPRSS2 cells the previous day at a concentration of 2.5 × 10^4^ cells/well. After 1 hour, the antibody-virus inoculum was removed and 0.85% methylcellulose in 2% FBS containing DMEM was overlaid onto the cell monolayer. Cells were incubated at 37°C for 16-40 hours. Cells were washed three times with 1X PBS (Corning Cellgro) and fixed with 125 μl of 2% paraformaldehyde in PBS (Electron Microscopy Sciences) for 30 minutes. Following fixation, plates were washed twice with PBS and 100 μl of permeabilization buffer, was added to the fixed cells for 20 minutes. Cells were incubated with an anti-SARS-CoV spike primary antibody directly conjugated with alexaflour-647 (CR3022- AF647) for up to 4 hours at room temperature. Plates were then washed twice with 1x PBS and imaged on an ELISPOT reader (CTL Analyzer). Foci were counted using Viridot (counted first under the “green light” set followed by background subtraction under the “red light” setting). IC50 titers were calculated by non-linear regression analysis using the 4PL sigmoidal dose curve equation on Prism 9 (Graphpad Software). Neutralization titers were calculated as 100% x [1- (average foci in duplicate wells incubated with the specimen) ÷ (average number of foci in the duplicate wells incubated at the highest dilution of the respective specimen).

### Immunogenetic analyses of antibody genes

The plasmid sequences were verified by Sanger sequencing (Macrogen sequencing, South Korea). The immunogenetic analysis of both heavy chain and light chain germline assignment, framework region annotation, determination of somatic hypermutation (SHM) levels (nucleotides) and CDR loop lengths (amino acids) was performed with the aid of IMGT/HighV-QUEST (www.imgt.org/HighV-QUEST) (26).

### Expression of human monoclonal antibodies

All transfections were done as described earlier (14). Briefly, expi293F cells were transfected with antibody expression plasmids at a density of 2.5 million cells per/ml using 1 mg/ml PEI-Max transfection reagent (Polysciences). Supernatants were harvested 4-5 days post-transfection and tested for their SARS-CoV-2 RBD binding potential by enzyme-linked immunosorbent assay (ELISA). Supernatant with positive RBD binding signals was next purified using Protein A/G beads (Thermo Scientific), concentrated using a 30 kDa or 100 kDa cut-off concentrator (Vivaspin, Sartorius) and stored at 4°C for further use.

### Electrochemiluminescence antibody binding assay

Binding analysis of SARS-CoV-2 mAb to spike protein was performed using an electrochemiluminescence assay as described earlier (14). Briefly, V-PLEX COVID-19 Panel 24 (Meso Scale Discovery) was used to measure the IgG1 mAb binding to SARS-CoV-2 spike antigens following the manufacturer’s recommendations. antigen coated plates were blocked with 150 μl/well of 5% BSA in PBS for 30 minutes. Plates were washed 3x with 150 μl/well of PBS with 0.05% Tween between each incubation step. mAbs were serially diluted for concentrations ranging from 10 μg/ml to 0.1 pg/ml and 50 μl/well were added to the plate and incubated for two hours at room temperature with shaking at 700rpm. mAb antibody binding was then detected with 50 μl/well of MSD SULFO-TAG anti-human IgG antibody (diluted 1:200) incubated for one hour at room temperature with shaking at 700rpm. 150 μl/well of MSD Gold Read Buffer B was then added to each plate immediately before reading on an MSD QuickPlex plate reader.

### Octet BLI analysis

Octet biolayer interferometry (BLI) was performed using an Octet Red96 instrument (ForteBio, Inc.) as described earlier (14). A 5 μg/ml concentration of each mAb was captured on a protein A sensor and its binding kinetics were tested with serial 2-fold diluted RBD (600 nM to 37.5 nM) and spike hexapro protein (100 nM to 6.25 nM). The baseline was obtained by measurements taken for 60 s in BLI buffer (1x PBS and 0.05% Tween-20), and then, the sensors were subjected to association phase immersion for 300 s in wells containing serial dilutions of RBD or trimeric spike hexapro protein. Then, the sensors were immersed in BLI buffer for as long as 600 s to measure the dissociation phase. The mean Kon, Koff and apparent KD values of the mAbs binding affinities for RBD and spike hexapro were calculated from all the binding curves based on their global fit to a 1:1 Langmuir binding model using Octet software version 12.0.

### Spike protein expression and purification

SARS-CoV-2 Spike-6P trimer protein carrying WA.1 was expressed and purified by transfecting expi293F cells using WA.1-spike-6P plasmids as described previously (14). Transfections were performed as per the manufacturer’s protocol (Thermo Fisher). Briefly, expi293F cells (2.5×10^6^ cells/ml) were transfected using ExpiFectamine™ 293 transfection reagent (ThermoFisher, cat. no. A14524). The cells were harvested 4-5 days post-transfection. The spike protein was purified using His-Pur Ni-NTA affinity purification method. Column was washed with Buffer containing 25 mM Imidazole, 6.7 mM NaH2PO4.H2O and 300 mM NaCl in PBS followed by spike protein elution in elution buffer containing 235 mM Imidazole, 6.7 mM NaH2PO4.H2O and 300 mM NaCl in PBS. Eluted protein was dialyzed against PBS and concentrated. The concentrated protein was loaded onto a Superose-6 Increase 10/300 column and protein eluted as trimeric spike collected. Protein quality was evaluated by SDS-PAGE and by Negative Stain-EM.

### Negative Stain – Electron Microscopy (NS-EM)

Spike protein was diluted to 0.05 mg/ml in PBS before grid preparation. A 3 μL drop of diluted protein (~0.025 mg/ml) was applied to previously glow-discharged, carbon-coated grids for ~60 sec, blotted and washed twice with water, stained with 0.75% uranyl formate, blotted and air-dried. Between 30-and 50 images were collected on a Talos L120C microscope (Thermo Fisher) at 73,000 magnification and 1.97 Å pixel size. Relion- 3.1 (27) or Cryosparc v3.3.2 (28) was used for particle picking and 2D classification.

### Sample preparation for Cryo-EM

SARS-CoV-2 spike-6P trimer incubated with the mAb (full-length IgG) at 0.7 mg/ml concentration. The complex was prepared at a 0.4 sub-molar ratio of mAb to prevent inter-spike crosslinking, mediated by bi-valent binding of intact antibody. The complex was incubated at room temperature for ~5 min before vitrification. Three μL of the complex was applied onto a freshly glow-discharged (PLECO easiGLOW) 400 mesh, 1.2/1.3 C-Flat grid (Electron Microscopy Sciences). After 20 s of incubation, grids were blotted for 3 s at 0 blot force and vitrified using a Vitrobot IV (Thermo Fisher Scientific) under 22°C with 100% humidity.

### Cryo-EM data acquisition

Single-particle Cryo-EM data for WA.1 spike-IgG complexes of mAb 002-02, 002-13 and 034-32 were collected on a 300 kV Titan Krios transmission electron microscope (ThermoFisher Scientific) equipped with Gatan K3 direct electron detector behind 30 eV slit width energy filter. Multi-frame movies were collected at a pixel size of 1.0691 Å per pixel with a total dose of 63 e/Å^2^ at defocus range of -0.7 to -2.7 μm.

### Cryo-EM data analysis and model building

Cryo-EM movies were motion-corrected by Patch motion correction implemented in Cryosparc v3.3.1 (28). Motion-corrected micrographs were corrected for contrast transfer function using Cryosparc’s implementation of Patch CTF estimation. Micrographs with poor CTF fits were discarded using CTF fit resolution cutoff to ~6.0 Å. Particles were picked using a Blob picker, extracted and subjected to an iterative round of 2D classification. Particles belonging to the best 2D classes with secondary structure features were selected for heterogeneous 3D refinement to separate IgG bound Spike particles from non-IgG bound Spike particles. Particles belonging to the best IgG bound 3D class were refined in non-uniform 3D refinement with per particle CTF and higher-order aberration correction turned on. To further improve the resolution of the RBD-IgG binding interface a soft mask was created covering one RBD and interacting Fab region of IgG and refined locally in Cryosparc using Local Refinement on signal subtracted particles. All maps were density modified in Phenix (29) using Resolve Cryo-EM. The combined Focused Map tool in Phenix was used to integrate high resolution locally refined maps into an overall map. Additional data processing details are summarized in Figure **S3-S6 and Table S1-2**.

The initial spike models for WA.1 (PDB:7lrt)) as well as individual heavy and light chains of the Fab region of an IgG (generated with Alphafold) (30) were docked into combine focused Cryo-EM density maps using UCSF ChimeraX (31). The full Spike-mAb model was refined using rigid body refinement in Phenix, followed by refinement in Isolde (32). The final model was refined further in Phenix using real-space refinement. Glycans with visible density were modelled in Coot and validated by Privateer (33, 34). Model validation was performed using Molprobity (35). PDBePISA (36) was used to identify mAb-RBD interface residue, to calculate buried surface area and to identify polar interaction. Figures were prepared in ChimeraX (31) and PyMOL (37).

### Molecular dynamics simulation

Molecular dynamics simulations were carried out to understand the effect of RBD mutations on the mAb binding. MD simulations were carried using AMBER99SB force field as implemented in GROMACS 2019. The system was solvated with TIP3P water model and neutralized with salts ([NaCl] =0.15 M,). Electrostatics were calculated using the PME method [24] with a real space cut-off of 10 Å. Van der Waals interactions were modelled using Lennard-Jones 6℃12 potentials with a 14 Å cut-off. The temperature was maintained at 300 K using V-rescale; hydrogen bonds were constrained using the LINCS algorithm [25]. Energy minimization was carried out to reach a maximum force of no more than 10 kJ/mol using steepest descent algorithm. The time step in all molecular dynamics simulations was set to 2 fs. Prior to the production run, the minimized systems were equilibrated for 5ns with NVT and followed with NPT at 300 K.

To calculate the Binding energies for the wild and the mutants, 200 snapshots were extracted from the last 20 ns of the 80ns production run. The extracted 400 frames for the wild and the variant subjected to MM/PBSA calculations using the gmx_MMPBSA tool. Before executing the calculations using gmx MMPBSA, PBC conditions were removed from the GROMACS output trajectory and protein-mAB complex were indexed. The AMBER99SB force field5 was used to determine the internal term (Eint), van der Waals (EvdW), and electrostatic (Eele) energies. Whereas GB-Neck2 model (igb = 8) was used to estimate the polar component of the solvation energy (GGB), while the non-polar solvation free energy (GSA) was calculated using the equation: Δ*GSA = γ* · △*SAS* + *ß*. Here the values of *γ* and *β* are 0.0072 kcal·Å-2·mol·^1^ and 0.

### Data availability

Atomic coordinates and Cryo-EM maps for reported structures are deposited into the Protein Data Bank (PDB) and the Electron Microscopy Data Bank (EMDB) with accession codes PDB-7U0Q and EMD-26263 for WA.1 Spike-6P in complex with mAb 002-02, PDB- 7U0X and EMD-26267 for WA.1 Spike-6P in complex with mAb 002-013, PDB-7UOW and EMD-26656 for WA.1 Spike-6P in complex with mAb 034-32. Immunoglobulin sequences are available in GenBank under accession numbers ON882061 - ON882244. All data needed to evaluate the conclusions in the paper are present in the paper and/or the Supplementary Materials. For materials requests, please reach out to the corresponding author.

### Statistical/Data analysis

Statistical analysis was performed with Prism 9.0.

## Acknowledgements

We thank Vineet D. Menachery and Pei-Yong Shi (The University of Texas Medical Branch) for providing the SARS-CoV-2mNG for the neutralization assays; Jason McLellan (The University of Texas) for providing the SARS-CoV-2 hexapro spike expression plasmid; Vinay Gupta (BD Biosciences India) and Aditya Rathee (ICGEB-TACF facility) for single-cell sorting; Satendra Singh and Ajay Singh (ICGEB, New Delhi) for technical support. The Cryo-EM data sets on the Titan Krios were collected at the National Center for CryoEM Access and Training (NCCAT) and the Simons Electron Microscopy Center located at the New York Structural Biology Center, supported by the NIH Common Fund Transformative High Resolution Cryo-Electron Microscopy program (U24 GM129539,) and by grants from the Simons Foundation (SF349247) and NY State Assembly. We also thank the staff of Robert P. Apkarian Integrated Electron Microscopy Core (IEMC) at Emory University, Atlanta for their support with preliminary sample screening on Talos Arctica.

## Author contributions

Experimental work, data acquisition and analysis of data by A.P., S.K., L.L., C.R.C., R.V., E.S.R., K.V.G., D.R.R., P.B., V.V.E., M.E.D.G., K.D., P.S., G.M., F.F., N.C., H.V., A.S.N., J.D.R., C.W.D., J.W., M.S.S., and E.A.O. Conceptualization and implementation by S.K., A.P., E.O., M.S.S., A.S., R.A., M.K.K., A.C. Manuscript writing by S.K., A.P., E.A.O., A.C., and M.K.K. All authors contributed to reviewing and editing the manuscript.

