## supplement information for "Molecular basis of SARS-CoV-2 Omicron variant evasion from shared neutralizing antibody response"

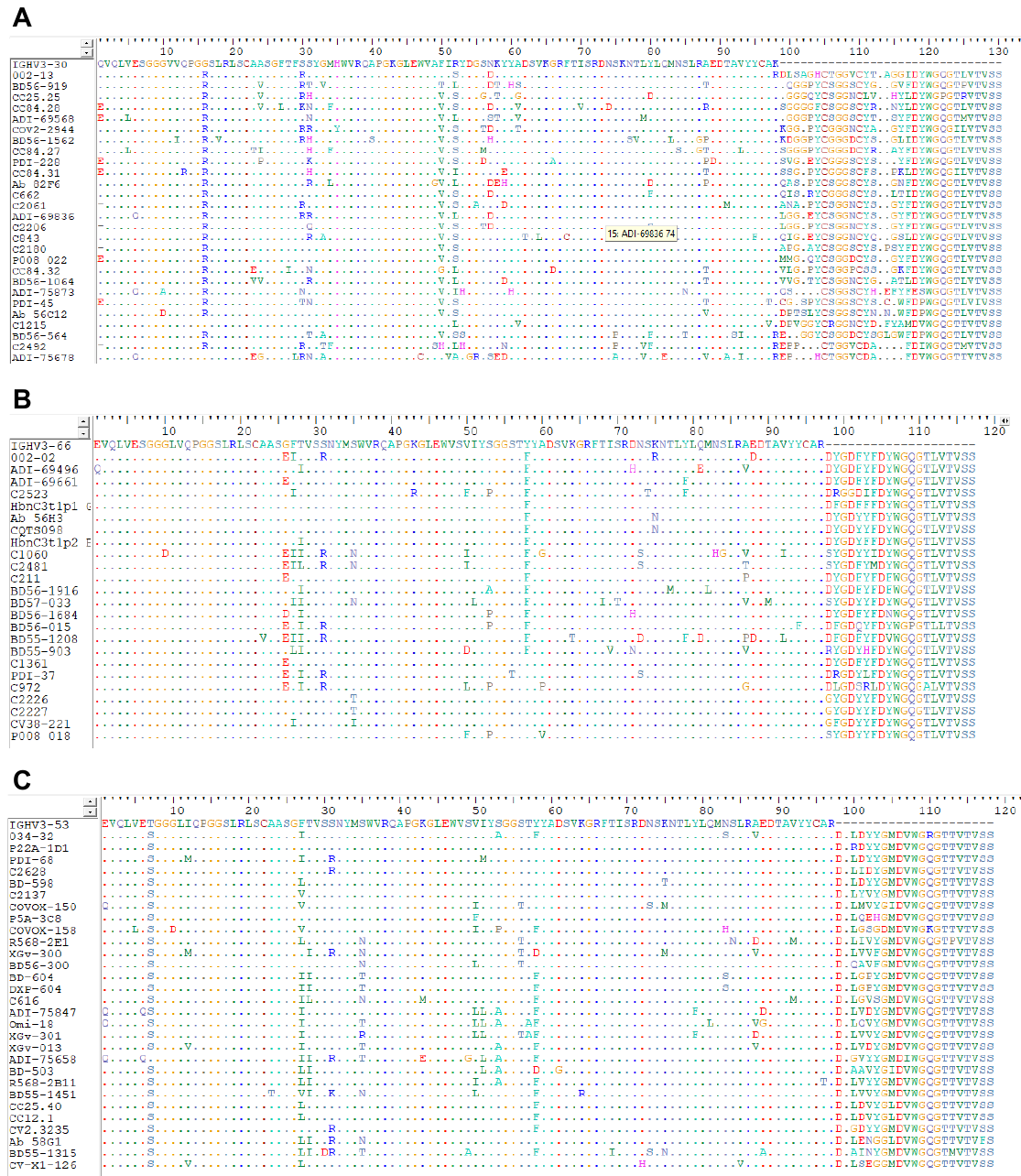

**Figure S1: Multiple sequence alignment of 002-13, 002-02 and 034-32 like shared mAbs.** (A-C) 002-13, 02-02 and 034-32 like mAbs with selected 002-13, 002-02 and 034-32 like IgH sequences encoded by IGHV3-30, IGHV3-66 and IGHV3-53 gene, respectively, identified from the COVID-19 patients and vaccinated individual. The respective germline gene sequence were used as the reference.

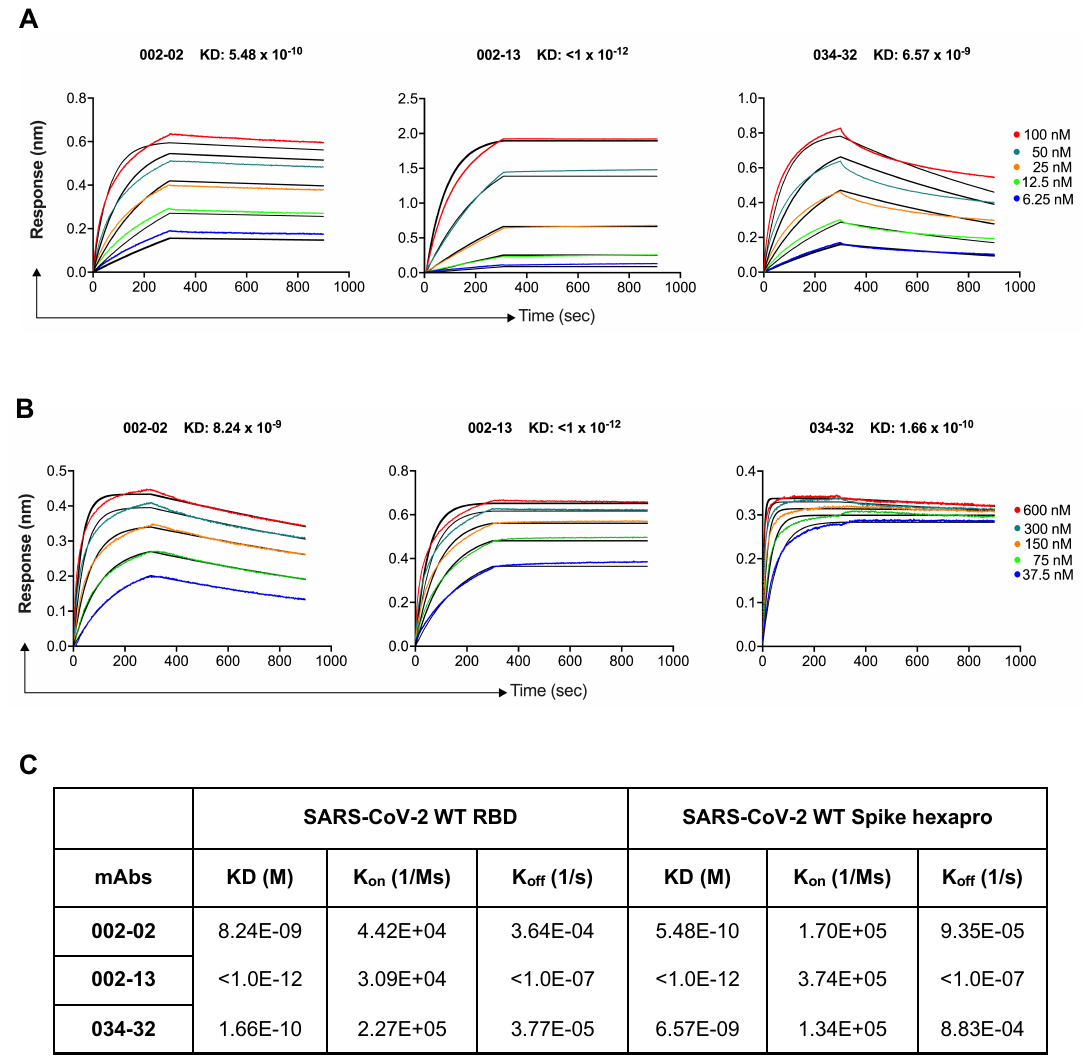

**Figure S2: Octet BLI sensor grams showing the Spike-6p binding affinities of the three potent mAbs, Related to Figure 1. (A)** In these assays, each mAb (5 ug/ml concentration) was captured on protein A sensors and its binding kinetics was tested with serial 2-fold diluted Spike hexapro protein **(A)** and RBD **(B)** (100 nM to 6.25 nM). Association was measured for 300 seconds followed by dissociation measurement for 600 seconds. **(B)** Describing the KD (M), K_on_ (1/Ms) and K_off_ (1/s) values of the four potent mAbs with RBD and Spike hexapro proteins.

**Figure S3. CryoEM data analysis and validation for WA.1 Spike-6P and 002-13 complex, Related to Figure 2. (A)** Representative electron micrograph. **(B)** Representative 2D-class averages. **(C)** Classification scheme and refinement that yielded final cryoEM map reconstruction. Boxed classes were selected for further processing and refinement. Boxed region containing one RBD complexed with one Fab in refined map masked for local refinement. **(D)** Gold standard Fourier shell correlation (FSC) curve of final overall (left) and locally refined (right) maps and resolution estimation based on 0.143 Fourier shell correlation criteria as indicated by blue line.

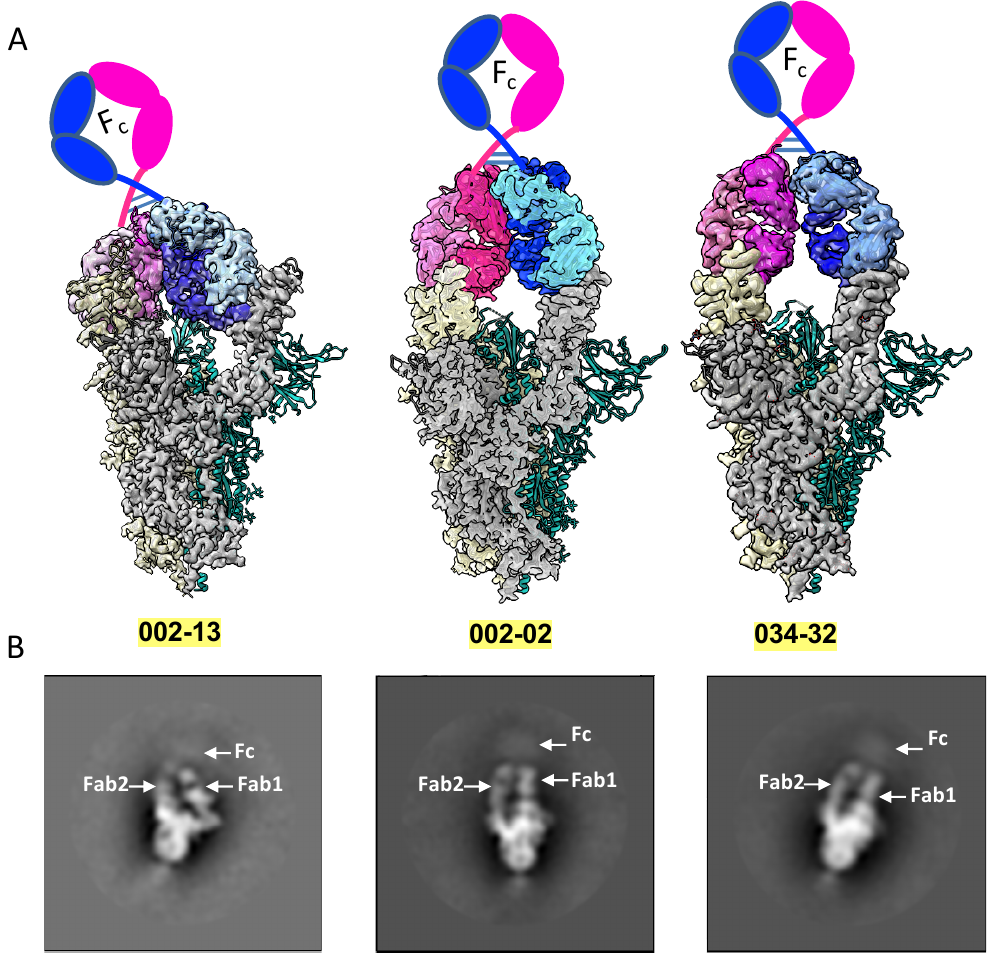

**Figure S4. Intra-spike bivalent IgG binding, Related to Figure 2, 3 and 4. (A)** Structural illustration of bivalent binding mAbs. Each protomer involve in intra-spike bivalent binding is shown in gray and yellow in spike trimer. Two RBDs on one spike trimer are occupied by two Fabs from the same IgG. **(B)** 2D-class averages derived from NS-EM data show the fuzzy Fc density supporting the intra-spike bivalent binding by two Fabs from same IgG.

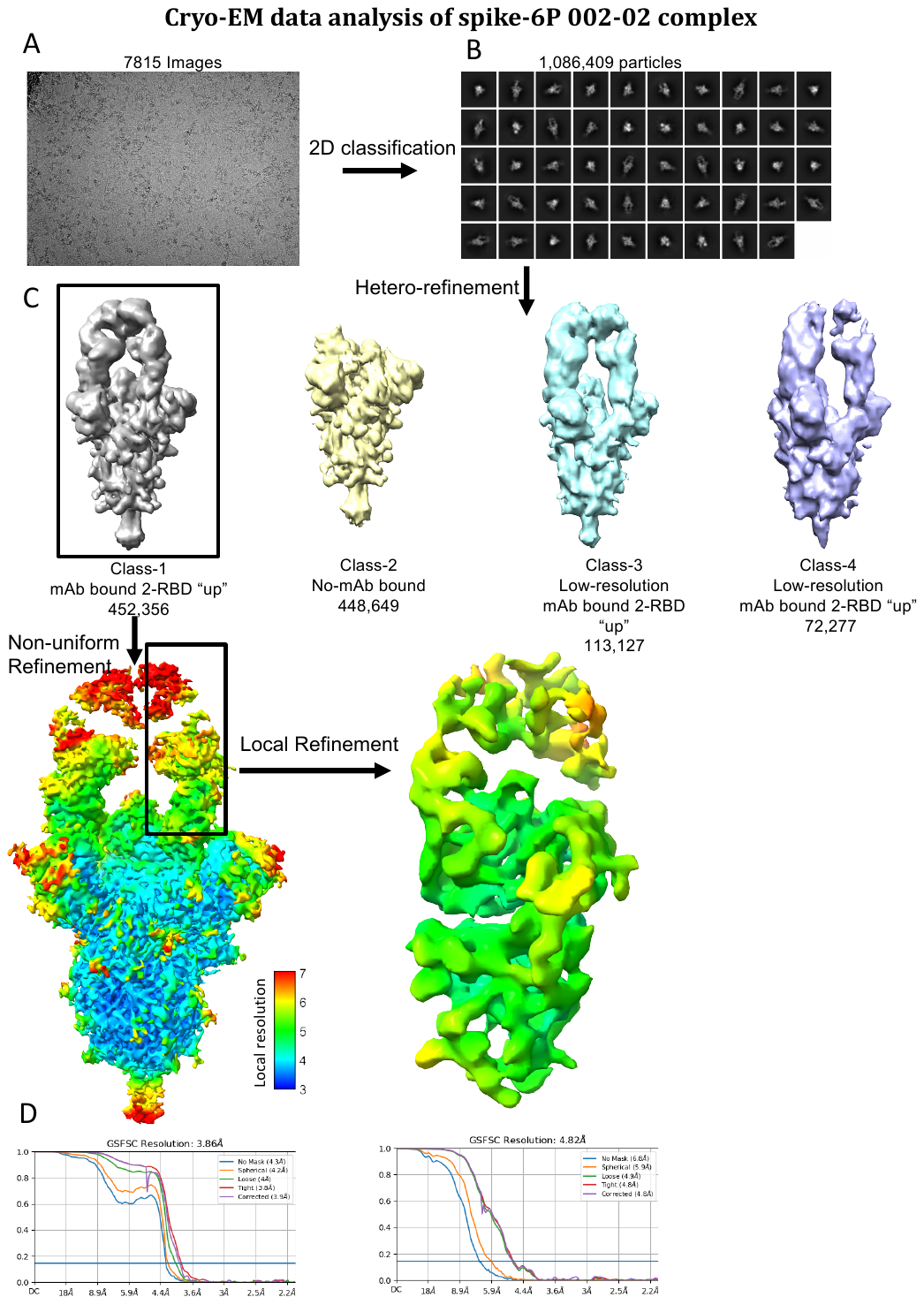

**Figure S5. Cryo-EM data analysis and validation for WA.1 Spike-6P and 002-02 complex, Related to Figure 3. (A)** Representative electron micrograph. **(B)** Representative 2D-class averages. **(C)** Classification scheme for 3D-classification and refinement that yielded final cryo-EM map reconstruction. Boxed classes were selected for further processing and refinement. Boxed region containing one RBD complexed with one Fab in refined map masked for local refinement. **(D)** Gold standard Fourier shell correlation (FSC) curve of final overall (left) and locally refined (right) maps and resolution estimation based on 0.143 Fourier shell correlation criteria as indicated by blue line.

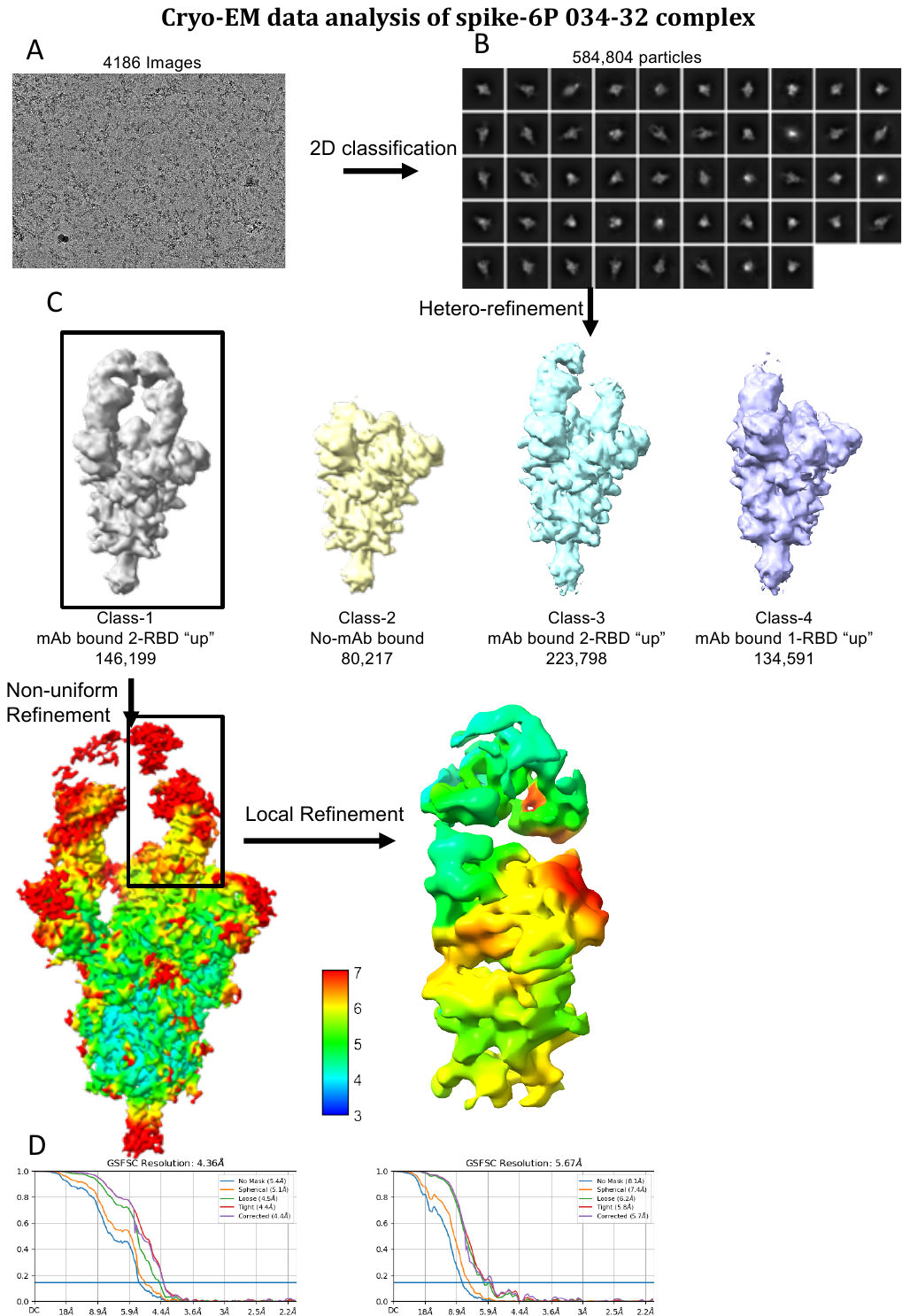

**Figure S6. Cryo-EM data analysis and validation for WA.1 Spike-6P and 034-32 complex, Related to Figure 4. (A)** Representative electron micrograph. **(B)** Representative 2D-class averages. **(C)** Classification scheme and refinement that yielded final cryo-EM map reconstruction. Boxed classes were selected for further processing and refinement. Boxed region containing one RBD complexed with one Fab in refined map masked for local refinement. **(D)** Gold standard Fourier shell correlation (FSC) curve of final overall (left) and locally refined (right) maps and resolution estimation based on 0.143 Fourier shell correlation criteria as indicated by blue line.

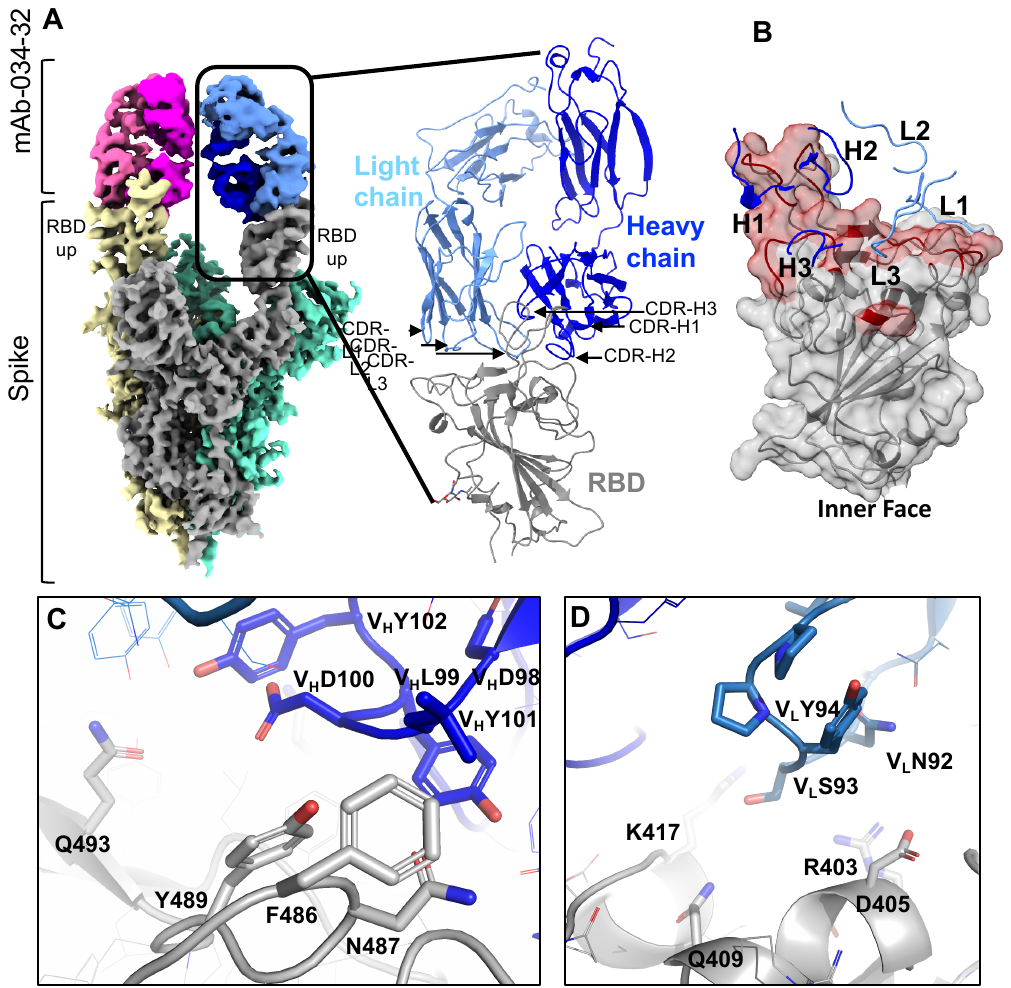

**Figure S7. Cryo-EM structure of 034-32 in complex with WA.1 spike trimer. (A)** Overall density map at contour level of 4.5 𝜎 showing the antibody binding two RBDs in the “up” conformation. Each protomer of Spike is shown in gray, yellow or green; light and heavy chains of each Fab region are shown in blue/ magenta and light blue/ pink, respectively. A model for one Fab-RBD complex is shown to the right; the positions of all Fab CDR regions are labelled. **(B)** Surface representation of RBD with the relative positions of all CDR loops. The mapped epitope surface in RBD is highlighted in maroon. **(C-D)** Interaction details of 034-32 CDR3 heavy **(C)** and light **(D)** chain loops with RBD.

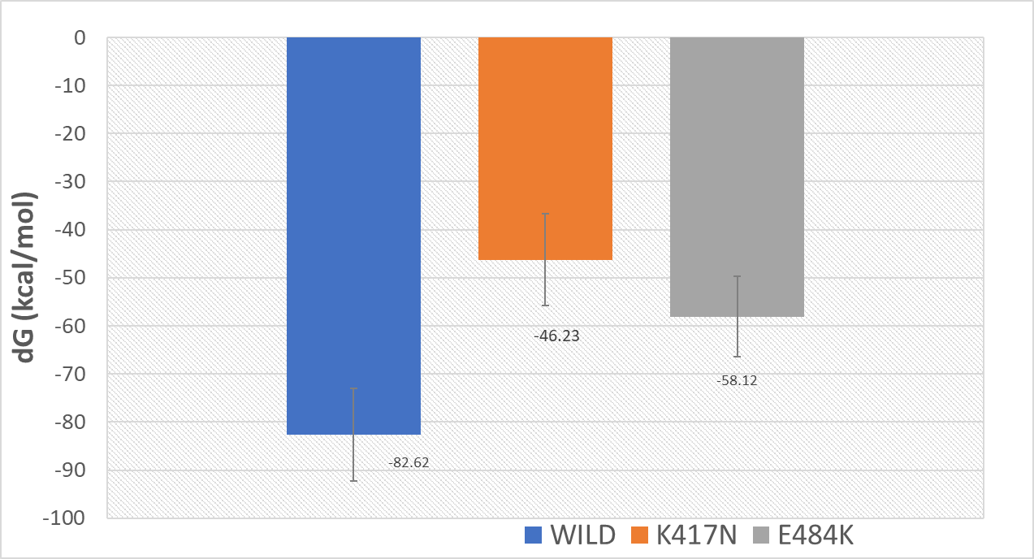

**Figure S8. Binding free energy of the WA.1 spike protein, K417N and E484K mutation.**

**Table S1: CryoEM data collection.**

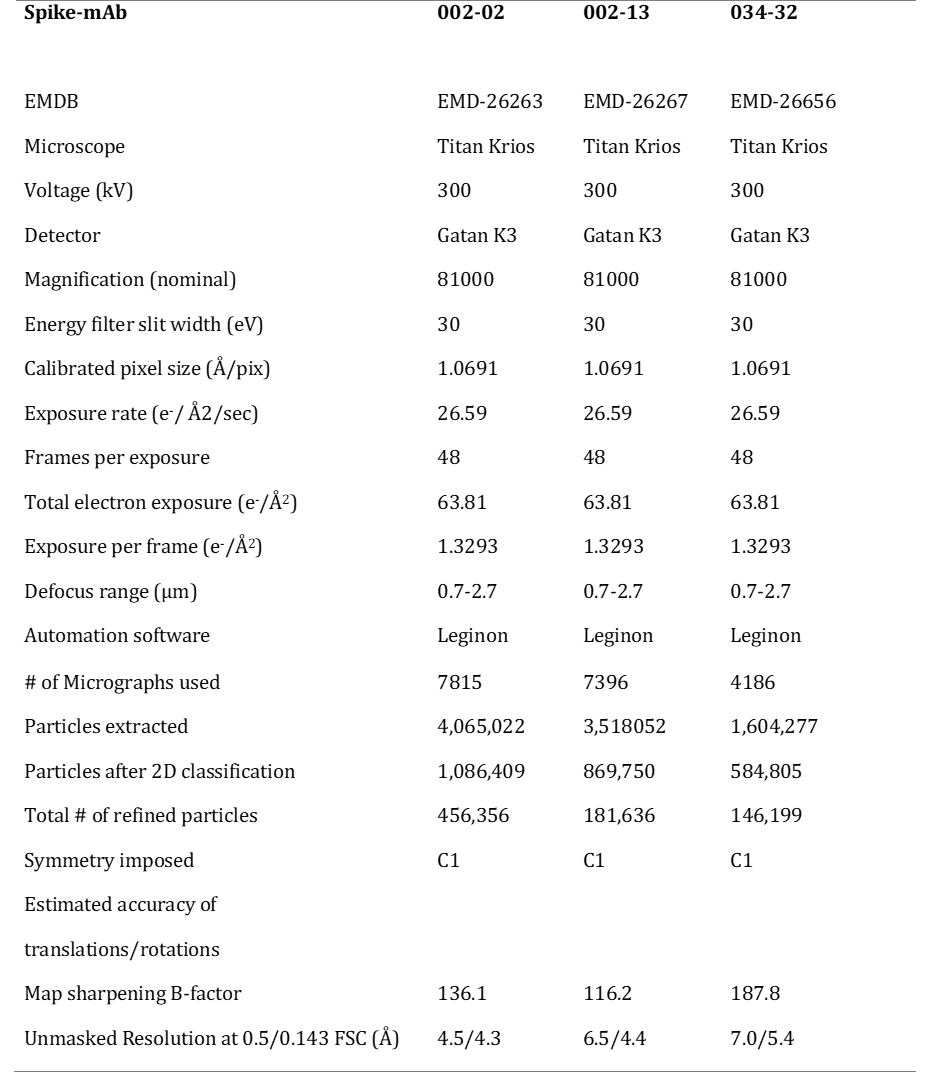

**Table S2: Model refinement and validation statistics.**

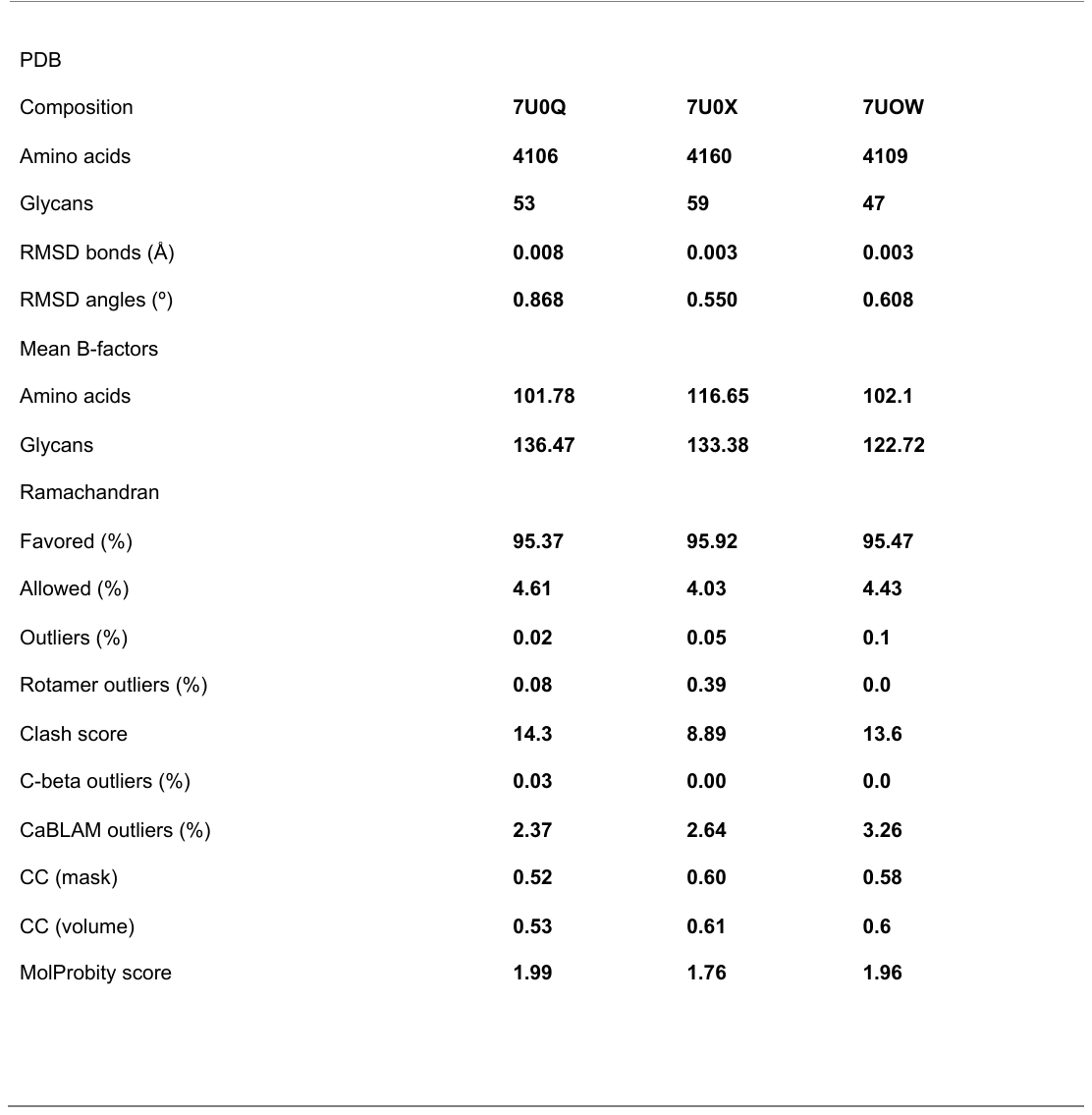
